# barbieQ: An R software package for analysing barcode count data from clonal tracking experiments

**DOI:** 10.1101/2025.09.11.675529

**Authors:** Liyang Fei, Jovana Maksimovic, Alicia Oshlack

**Author notes:** These authors jointly supervised this work (Jovana Maksimovic, Alicia Oshlack).

## Abstract

**Motivation:** A “clone” encompasses a progenitor cell and its progeny cells. Tracking clonal composition as cells differentiate or evolve is useful in many fields. Various single-cell lineage tracing (clonal tracking) technologies use unique DNA barcodes that are passed from progenitor cells to their offspring. The barcode count for each sample indicates cell number in clones. However, analysis of barcode count data is often bespoke and relies on visualisations and heuristics. A generalized workflow for preprocessing and robust statistical analysis of barcode count data across protocols is needed.

**Results:** We introduce *barbieQ*, a Bioconductor R package for analysing barcode count data across groups of samples. It provides data-driven quality control and filtering, extensive visualisations, and two statistical tests: 1) Differential barcode proportion (differences in proportions between sample groups), and 2) Differential barcode occurrence (differences in presence/absence odds between groups). Both tests handle complex experimental designs using regression models and rigorously account for sample-to-sample variability. We validated both tests on semi-simulated, real data and a case study, demonstrating that they hold their size, are sufficiently powered to detect true differences, and outperform existing approaches.

## Introduction

Cellular DNA barcoding is a powerful cell clonal tracking (lineage tracing) technique also referred to as lineage tracing. It incorporates unique DNA barcodes into a cell’s genome, thereby genetically labelling each cell in a population.^1^ The DNA barcodes are inherited by daughter cells, allowing the origin of each progeny cell to be identified. Collectively, an initial cell and all its progeny are referred to as a “clone” or “lineage”. Tracking clonal behaviour reveals underlying clonal heterogeneity, leading to widespread use of DNA barcoding in studies of haematopoietic stem cell differentiation, tumour cell evolution, and embryonic development.^2–7^ While each barcode serves as a unique clonal identifier, the total number of cells in each clone can be determined by sequencing and counting the barcode frequencies, thereby estimating relative clonal abundance. Based on the observed clonal abundance patterns, many studies then investigate the features of the initial cells, such as gene expression, gene activity and epigenomic patterns, that relate to clonal outcomes.^8–11^

Cellular genetic barcoding involves several steps, including barcode sequence design, barcode incorporation into cells, and barcode sequencing, each supported by several published protocols.^12–24^ These protocols have inspired a series of bespoke workflows, protocol-specific bioinformatics tools, and generalised software for barcode sequence recovery.^25–32^ Typically, next-generation-sequencing (NGS) combined with barcode-specific PCR amplification is used to efficiently capture barcode-specific DNA from bulk cell samples. The *CellBarcodeSim* software can simulate barcode data, including the technical effects introduced throughout the barcoding processes.^32,33^

Genetic barcoding-based studies can be broadly categorized into two analytical aims: lineage tracing and clonal analysis. Although often used interchangeably, we distinguish them here by analytical focus and data input. Lineage tracing reconstructs hierarchical lineage trees by resolving progeny relationships across successive divisions, often coupling clonal and transcriptomic information to infer lineage trajectories.^34,35^ In contrast, clonal analysis treats all progeny from a common ancestor as a clone and focuses on summarizing progenitor fate, typically by quantifying cell type distribution within each clone, where molecular measurements are often limited to the progenitor state.^36,37^

In this work, we focus on clonal analysis, for which lentiviral static barcoding is particularly suited as it provides large-scale barcode libraries with defined, static sequences. This framework suits experimental systems such as *in vitro* cultures or *in vivo* regenerative models, where cells are harvested, labelled, transplanted, and re-sampled across conditions or as parallel transplants. A key biological assumption underlying clonal analysis is that every cell carrying a barcode, including all its progeny inherited that barcode from a single originally labelled progenitor. This means the number of cells per clone can be quantified by counting a certain barcode sequence. In these settings, each progenitor cell is assumed to carry a unique barcode, and each sample comprises progeny from multiple progenitors under a given condition, providing a basis for comparing clonal contributions across conditions. Regardless of extraction and alignment approach, most pipelines output a matrix of barcode read counts across samples.

A barcode matrix typically contains thousands of barcodes across tens to hundreds of samples. The overall read count for a single barcode represents the number of cells derived from the originally barcoded cell. However, in studies such as *in vivo* haematopoietic stem cell tracking, not all progeny cells can be collected, so barcode counts reflect the relative abundance of each lineage in a sample. This contrasts with studies such as *in vitro* tumour heterogeneity tracking, where all progeny cells can be collected and sequenced, allowing for barcode counts to be interpreted as absolute abundance. However, even in the second scenario, researchers may not sequence all of progeny cells collected or know the exact total cell count. Consequently, it is more common and practical to interpret barcode count as relative or proportional abundance.

Several bioinformatics tools that offer a range of visualization approaches and heuristic interpretation strategies have been developed to analyse barcode count matrixes. *barcodetrackR* on Bioconductor, and *CellDestiny* and *bartools* on GitHub^28–31,38–41^ are widely used and share some common features. All three tools typically use heatmaps to visualise the abundance of each barcode per sample. *BarcodetrackR* and *bartools* display the log normalised barcode proportion or counts per million (CPM) for selected barcodes (e.g. top 10 most abundant per sample), whilst *CellDestiny* displays the arcsine transformed proportion across all barcodes. Additionally, *CellDestiny* provides line plots and bubble plots, whilst *bartools* provides stacked bar plots and bubble plots for displaying barcode abundance across multiple samples.

Furthermore, all three tools also summarise sample characteristics using the per-sample Shannon diversity index and number of features (detected unique barcodes per sample). For assessing sample differences, *barcodetrackR* uses MDS plots and *bartools* uses PCA plots. Both assess sample similarities by displaying sample pair-wise correlations using the Pearson correlation coefficient. Notably, *CellDestiny* characterises the distribution of barcode abundance within each sample by visualising the cumulative barcode proportion per sample.

However, the three tools differ in how they characterize barcode changes between sample groups. *barcodetrackR* calculates the log fold change (LFC) of barcode proportions between two groups of samples and categorizes barcodes using a heuristic threshold. *CellDestiny* compares relative barcode abundance across groups by first applying column-wise and then row-wise normalisation to the barcode count matrix, then classifying the barcodes as either group-specific or universal based on whether each group passes a heuristic threshold. Neither of these tools provides a statistical test. *bartools* tests the odds of barcode presence/absence between two groups of progeny cells using a hypergeometric test, typically applied in the context of two clusters in single cell barcoding data.

Lineage tracing experiments often involve complex experimental designs, including multiple time points, genotypes, treatments, and both biological and technical replicates. For example, in a xenograft-based haematopoiesis study, the clonal output was determined by a combination of different cell types, tissues and time points from which cells were collected.^28,42^ Despite this complexity, existing tools lack statistical frameworks capable of appropriately testing for differences between groups of samples. Specific limitations include,

1. The LFC approach, the relative proportion approach or the hypergeometric test cannot directly handle multiple conditions within a single factor.
2. None of the approaches can robustly address within-group variability. Technical replicates can be aggregated by *CellDestiny* and *bartools*, however, they cannot rigorously address the differences within replicates and between samples.
3. Both LFC and relative proportion approaches rely on heuristic thresholds to characterise differences in barcode abundance between samples, which can result in subjective and unreproducible interpretations.

As such, there are a lack of robust and rigorously validated statistical testing models specifically designed for population based (bulk) barcode count data that can adequately address barcode variability between conditions. A study associated with the DEBRA approach has explored adapting the *DESeq, DESeq2*, and *edgeR* models for barcode count data, however, it lacks thorough validation and comprehensive assessment of the performance of these models.^43–45^

This paper presents the *barbieQ* R Bioconductor package, which provides barcode normalization, data-driven filtering, rich visualisations and statistical testing accommodating the multifactorial designs that are common in the lineage tracing field to enable reproducible analysis.^46^ The statistical tests in *barbieQ* build upon a linear regression framework to account for variation in barcodes arising from various sources and can robustly detect significant changes in both barcode abundance and barcode presence/absence between groups of samples. We have used four publicly available datasets to demonstrate and validate *barbieQ*. We show that *barbieQ* controls false discovery rate using both real and semi-simulated datasets and demonstrate its ability to detect real barcode differences in public data.

## Results

### Package overview

The *barbieQ* package workflow consists of data preprocessing and statistical testing, supported by corresponding visualisations. Briefly, preprocessing loads in the barcode counts and produces a “*barbieQ*” object, preparing data for efficient and accurate interpretation. Statistical testing of the preprocessed “*barbieQ*” object identifies barcodes with significant changes between sample groups in either occurrence or proportion. These results can be visualised using dedicated functions. (Figure 1)

**Figure 1.**
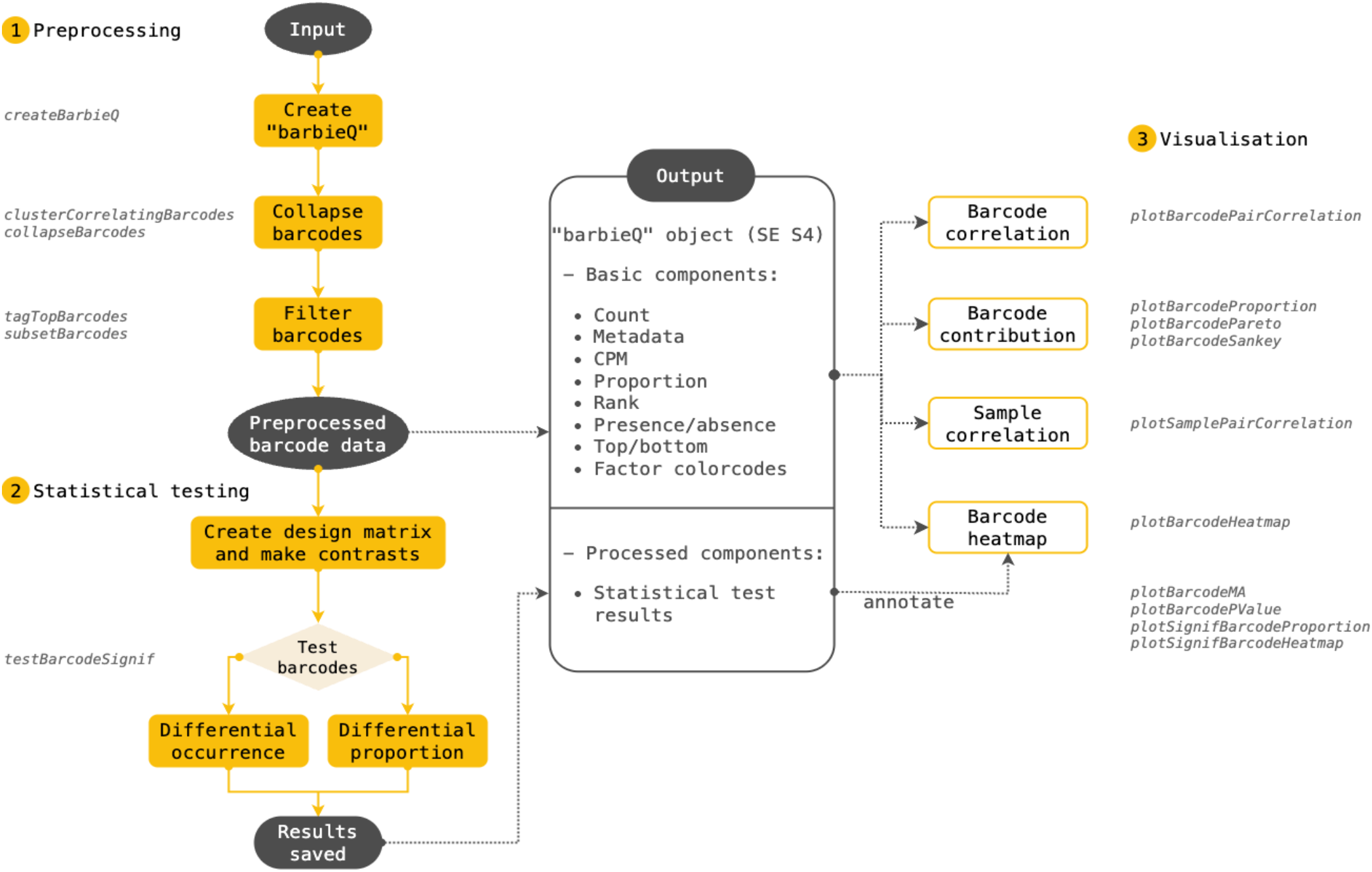
Workflow of *barbieQ* package. *barbieQ* consists of three modules: preprocessing, statistical testing, and visualisation. 1) Preprocessing starts by creating a “*barbieQ*” object, and updates it by collapsing highly correlated barcodes, and filtering out lowly abundant barcodes, ensuring an accurate and clean set of barcodes for subsequent interpretation and visualization. Dedicated functions for each step are listed. 2) Statistical testing applies to the preprocessed “*barbieQ*” object with choices for differential occurrence or differential proportion test, and stores the results in the object, which can be visualised. 3) Dedicated visualisations are available at any stage.

More specifically, preprocessing starts by using `“createBarbieQ”` to read in a *matrix* of the barcode counts, optionally including a *data*.*frame* with sample metadata, which initializes a “*barbieQ*” object that is updated throughout the workflow. The “*barbieQ*” object is built upon the *SummarizedExperiment* structure in *R*, consisting of assays storing raw and processed count matrices, barcode-oriented metadata, and sample-oriented metadata. Based on the imported barcode count matrix, the “`createBarbieQ`” function calculates the barcode counts per million (CPM), proportion and ranking per sample, as well as the presence/absence (occurrence), and saves the results for each metric as a separate matrix in assays. The definitions of those metrics are listed in Table 2. The function also calculates the diversity index, number of unique barcodes (n_barcode) and number of total barcode counts (n_count) for each sample and saves those sample features as part of the sample data (column data), along with any imported sample metadata. Barcode occurrence is, by default, defined using a CPM threshold of 1, corresponding approximately to a single read in datasets with a library size of ∼1 million. CPM is used rather than raw counts to account for differences in library size across samples. Since barcode detection, particularly with lentiviral static barcoding, relies on strict alignment to a known reference library, spurious calls are unlikely. Nevertheless, users are encouraged to select a threshold appropriate for their library size and experimental context.

**Table 1.**
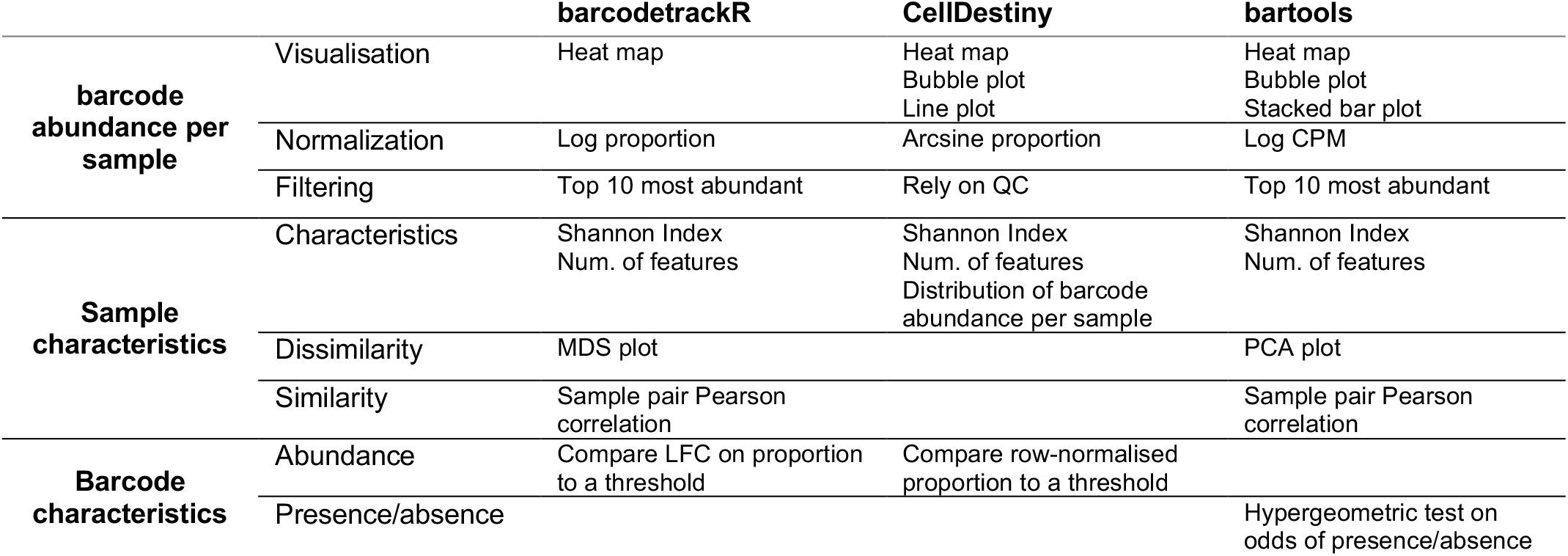
Feature comparison of existing tools for barcode count analysis.

**Table 2.** *barbieQ* object structure and components.

| Type | Component | Definition | Initiated by Function |
| --- | --- | --- | --- |
| **Assay** | Count | Barcode count per sample | *createBarbieQ* |
|  | CPM | Barcode count / total barcode count per sample * 1M | *createBarbieQ* |
|  | Proportion | Barcode count / total barcode count per sample | *createBarbieQ* |
|  | Occurrence | Presence/absence: barcode CPM >=1 or CPM <1 | *createBarbieQ* |
|  | Rank | Rank of barcode count per sample | *createBarbieQ* |
|  | isTop | Top/bottom: barcode being main contributor or low count per sample | *tagTopBarcodes* |
| **Sample data**<br>**(columnData)** | SampleMetadata | Sample experimental conditions (factors) | *createBarbieQ* |
|  | N_barcode | Number of detected barcodes (total occurrence) per sample | *createBarbieQ* |
|  | N_count | Number of total barcode counts (library size) per sample | *createBarbieQ* |
|  | Diversity index | Sample Shannon diversity index | *createBarbieQ* |
|  | Cluster | Barcode cluster of high correlation | *clusterCorrelatingBarcodes* |
| **Barcode data**<br>(rowData) | isTop | Barcode determined as top/bottom considering all samples | tagTopBarcodes |
|  | Significance | Significance of barcode abundance changes between sample groups | testBarcodeSignif |

`Next, “clusterCorrelatingBarcodes”` and `“collapseBarcodes”` can be used to detect and collapse any barcodes co-occurring in the same cell. `“plotBarcodePairCorrelation”` can be used to inspect the correlation between every pair of barcodes. Filtering barcodes with, `“tagTopBarcodes”` and `“subsetBarcodes”` tags barcodes as top/bottom contributors and filters out low-count/low-contribution barcodes. Visualizations using `“plotBarcodeProportion”`, `“plotBarcodePerato”` and `“plotBarcodeSankey”` can be used for inspecting each barcode’s contribution before and after filtering. Each step stores the results in the “*barbieQ*” object (Table 2), and recalculates assays such as CPM, proportion, rank, and occurrence when barcodes are collapsed or filtered.

The preprocessed “*barbieQ*” object, generated by collapsing barcodes within the same cell and removing low-abundance barcodes, provides a clean set of barcodes for statistical testing and allows accurate estimation of barcode proportions.

For statistical testing, `“testBarcodeSignif”` is applied to a “*barbieQ*” object and requires a user-defined design matrix indicating the sample conditions and other experimental factors, as well as user-defined contrasts that specify which sample groups to compare. Users can choose to test either differential occurrence or differential proportions of barcodes between the specified sample groups. The results are stored in barcode data in the “*barbieQ*” object (Table 2) and can be visualised by applying `“plotBarcodeMA”`, `“plotBarcodePValue”`, `“plotSignifBarcodeProportion”`, or `“plotSignifBarcodeHeatmap”` to the “*barbieQ*” object.

Additional visualizations, such as `“plotSamplePairCorrelation”`, `“plotBarcodeHeatmap”` and barcode PCA, which use barcode CPM, are available before or after statistical testing, to inspect sample heterogeneity and barcode variability.

### Barcode Preprocessing with barbieQ

We demonstrate the effectiveness of the implemented preprocessing functions in barbieQ by applying to the example dataset of 30 selected samples from the monkey HSPC data.

#### Collapsing highly correlated barcodes

The `“clusterCorrelatingBarcodes”` function is used to identify clusters of highly correlated barcodes, which are likely barcodes that have been integrated into the same initial cell and, therefore, co-exist in the progeny cells. This phenomenon has been previously observed and, if not properly accounted for, leads to overcounting of cells and distortion of proportions violating the assumption of one-to-one mapping of barcodes to cells.^31^ To address the issue, this function calculates the correlation of the proportions across samples of every pair of barcodes and classifies highly correlated pairs based on thresholds for correlation coefficient and mean proportion. The function provides several options for correlation methods and proportion transformation methods; by default, it uses Pearson correlation coefficient (*r*) of asin-sqrt transformed proportion. Highly correlated pairs are grouped into clusters containing two or more barcodes. Barcodes within a cluster are considered to co-exist in one clone and should be merged. `“collapseBarcodes”` selects the representative barcode as the one with the highest mean proportion to maintain a consistent barcode-clone association.

We applied this approach to the monkey HSPC data. Barcode pairs with a correlation coefficient greater than 0.95 and an average mean proportion exceeding 0.001 were classified as paired barcodes belonging to a cluster (Figure 2A). In total, 9 barcodes formed 4 clusters, each containing 2∼3 barcodes (Figure 2B). Barcodes within the same cluster exhibited similar proportion patterns across samples (Figure 2C). These barcode clusters were subsequently collapsed for downstream analysis, after which the previously excluded low-count barcodes were reintroduced for more accurate proportion estimation. These observations have been made in a published study, and unpublished data we analysed, where paired barcodes were validated through single-cell RNA sequencing of the initial cells, confirming the co-existence and co-expression of the DNA barcodes pairs (Supplementary Figure 1A).^31^ Preprocessing results for three additional datasets (AML data, HSPC xenograft data, and mixture data) can be found in Supplementary Figure 1.

**Figure 2.**
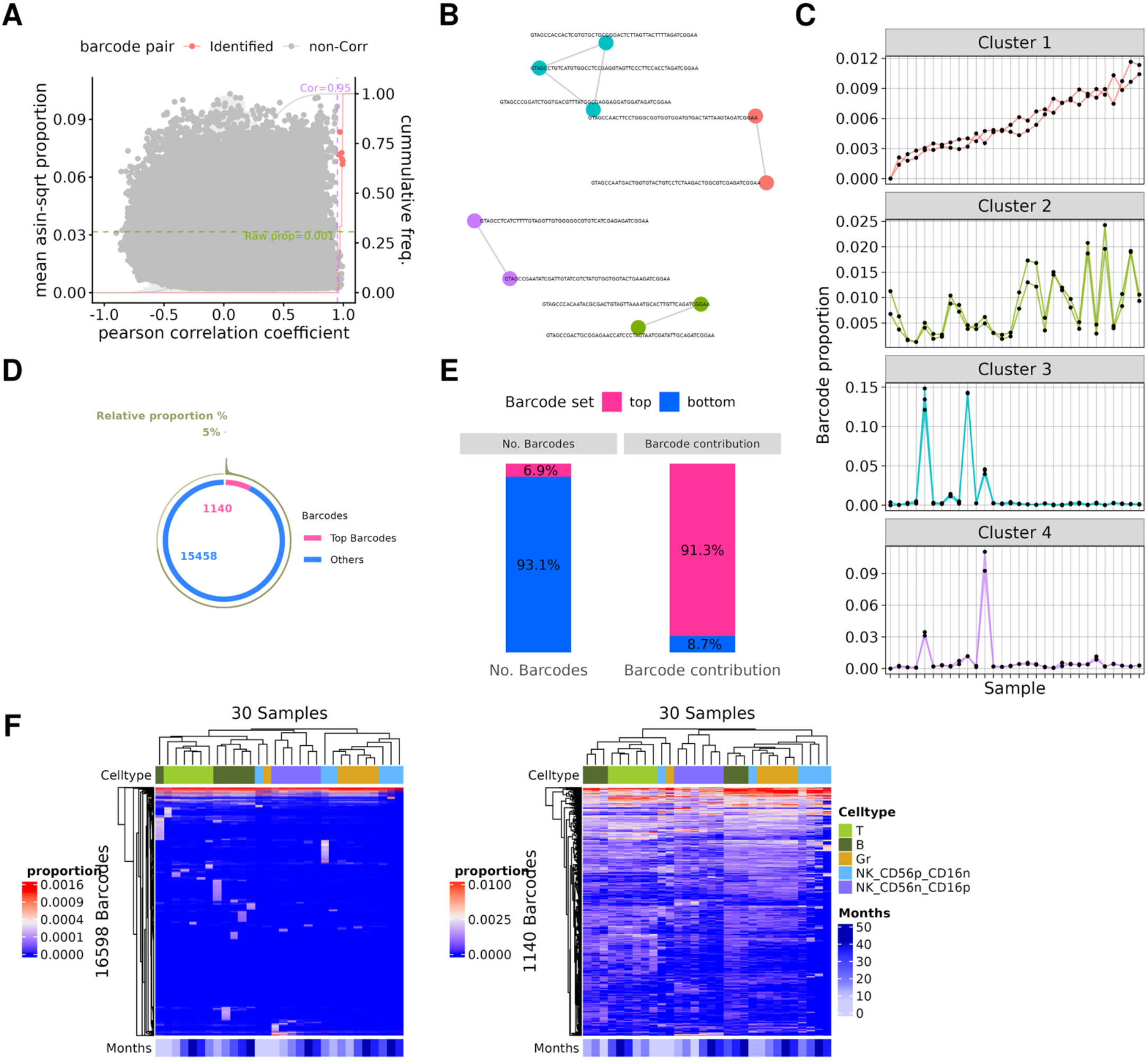
Preprocessing results on monkey HSPC data. (**A**) Mean proportion (asin-sqrt transformed) and Pearson correlation coefficients for each pair of barcodes out of the selected not-low-count barcodes using the `“plotBarcodePairCorrelation”` function; (**B**) Cluster of highly correlated barcodes visualised using the `“clusterCorrelatingBarcodes”` function; (**C**) Barcode proportion across samples within each cluster; each line represents an individual barcode; (**D**) Proportion of each barcode’s mean CPM out of all barcodes (outside circle), color-coded by “top” or “bottom” contributors as tagged by the filtering strategy (inside circle), using the `“plotBarcodePareto”` function; (**E**) Fraction of “top” or “bottom” contributor barcodes and their cumulative mean proportion, using the `“plotBarcodeSankey”` function; (**F**) Heatmap of barcodes on proportion before (**left**) and after (**right**) filtering as visualised using the `“plotBarcodeHeatmap”` function; heatmap colour scale mapped to asin-sqrt transformed proportion but labelled by raw proportion.

#### Barcode filtering

Barcode filtering is commonly used in barcode analysis pipelines to reduce the large number of minimally contributing barcodes that can affect statistical power and multiple testing burden. The `“tagTopBarcodes”` function uses a data-driven approach to classify barcodes as “top” and “bottom” contributors, where “top” barcodes that contribute substantially to the total counts are retained while “bottom” barcodes with minimal contributions are removed. Barcode proportions are calculated for each sample and ranked from highest to lowest. A cumulative proportion is then calculated at each rank, starting from the most abundant barcode. Barcodes ranked above the cumulative proportion cutoff (default = 0.99) are tagged as “top” in a sample, indicating they are the most abundant. Across the dataset, barcodes tagged as “top” in at least *n* samples (*n* being the minimum sample group size) are classified as “top” and retained across the whole dataset. This approach ensures that consistently low-abundance barcodes are filtered out while preserving those present in at least one sample group.

We applied this approach to the monkey HSPC dataset, using the default proportion cutoff of 0.99 and a minimum group of n = 6; this was chosen because the 30 samples feature 5 groups of cell types with a minimum group size of 6. This resulted in the classification of 1140 barcodes as “top” barcodes out of a total of 16598. The mean proportion of each barcode was visualised using the *barbieQ* function `“plotBarcodeProportion”` (Figure 2D). These “top” barcodes represent 6.9% of all barcodes accounting for 91.3% of total barcode counts in the dataset as shown in Figure 2E using the `“plotBarcodeSankey”` visualisation. By filtering out consistently low abundance “bottom” barcodes, we retained a focused, clean, and interpretable set of “top” barcodes (Figure 2F).

### Performance of *barbieQ* statistical testing

To assess differences in barcode counts between sample groups that represent clonal outcomes, *barbieQ* can analyse barcode changes in two ways: differential proportion and differential occurrence. Differential proportion identifies barcodes with significantly different proportions between the samples of two conditions, whilst differential occurrence identifies barcodes that are more frequently present in samples of one condition compared to another.

#### Model description and hypothesis

Both statistical tests are based on linear regression models that assess each barcode individually, while accounting for barcode variability between samples groups. A key feature of regression models is their ability to handle complex experimental designs using a design matrix that accounts for multiple factors and estimates the variance associated with each factor. This enables the tests to determine whether the factor of interest significantly changes with respect to barcode occurrence or proportion, while accounting for the effects of other confounding factors. To ensure that the correct hypothesis is tested (i.e., the factor or group of interest), the design matrix must be carefully specified. Guidance on constructing appropriate design matrices, including for complex experimental designs, has been provided by Law et al.^47^

The differential proportion test leverages the “*limma*” linear regression framework, modelling barcode proportion as a function of the factors included in the design matrix.^48,49^ Linear models assume that 1) residuals are approximately normally distributed and 2) residual variance is constant across levels of the independent variables (i.e., homoscedasticity). However, proportional data often violates these assumptions, particularly near the boundaries of 0 and 1. Inspired by Phipson et al., *barbieQ* differential proportion test models barcode proportions after applying an arcsine square root (asin-sqrt) or logit transformation to stabilize the variance across the range of proportions.^50^ Within this framework, the null hypothesis for each barcode is that the coefficient associated with the factor of interest in the design matrix is equal to zero, indicating no difference in transformed barcode proportions between groups after accounting for other covariates. Rejection of the null hypothesis therefore implies that the biological factor of interest is associated with a change in barcode proportion.

The differential occurrence test uses a logistic regression model, modelling the log-odds of barcode presence (i.e., log(probability of presence / probability of absence)) as a function of biological factors. The model is combined with Firth’s penalised likelihood to reduce bias in the maximum likelihood estimates of regression coefficients.^51,52^ This correction addresses inflated standard errors that can arise in extreme cases of complete or near-complete separation, which are more likely to occur in small sample size cases (e.g., n = 3), a scenario not uncommon in clonal tracking studies. Barcode presence or absence in each sample is treated as a binary outcome, and the model is specified using a design matrix in the same way as in the differential proportion test. Again, the null hypothesis for each barcode is that the coefficient associated with the factor of interest in the design matrix is equal to zero, indicating no difference in the probability of barcode occurrence between groups after adjusting for other covariates (equivalently, testing whether the odds ratio between groups differs from 1). Rejection of the null hypothesis therefore implies that the biological factor of interest is associated with a change in barcode occurrence.

The function `“testBarcodeSignif”` allows users to specify the statistical test and define the factor of interest through a design matrix with the desired contrast. The results of the test are saved to the *rowData* of the *barbieQ* object, aligned with the corresponding barcodes. They contain key statistics such as group-wise means and differences in proportion or odds ratios in occurrence, along with *P*-values adjusted for multiple testing using the Benjamin-Hochberg method by default.^53^ The test results can be visualised using the functions listed in Figure 1.

#### Type I error control under null scenario using real data

To evaluate the Type I error (false positives) of the statistical tests in *barbieQ*, we applied them to a variety of semi-simulated “null” datasets. From several datasets (mixture data, HSPC xenograft data, and AML data), we selected sets of samples with homogeneous conditions, where no biologically meaningful differences were expected. For each sample set, we generated 100 simulated null datasets by randomly assigning samples into two groups without replacement. We then applied the `“testBarcodeSignif”` function to perform differential occurrence and differential proportion tests, assessing the number of barcodes with significant changes (*P*-value < 0.05) between the two groups, which represent false positives. We also varied the number of samples per group to determine the effect of group size on Type I error rate (n = 3, 4, …, up to half the total number of the null samples).

Figure 3A shows the design of the Mixture dataset. Cells from Pool1 and Pool2 were counted and mixed in equal ratios to produce a mixed pool. Twelve baseline samples (each from the mixture of Pool 1 and Pool 2 in equal proportions) with varying total cell numbers (n = 20, 40, 80, 160, 330, and 660k, two replicates each) were used for Type 1 error evaluation as relative barcode abundance should be consistent regardless of the technical effects related to total cell number.

**Figure 3.**
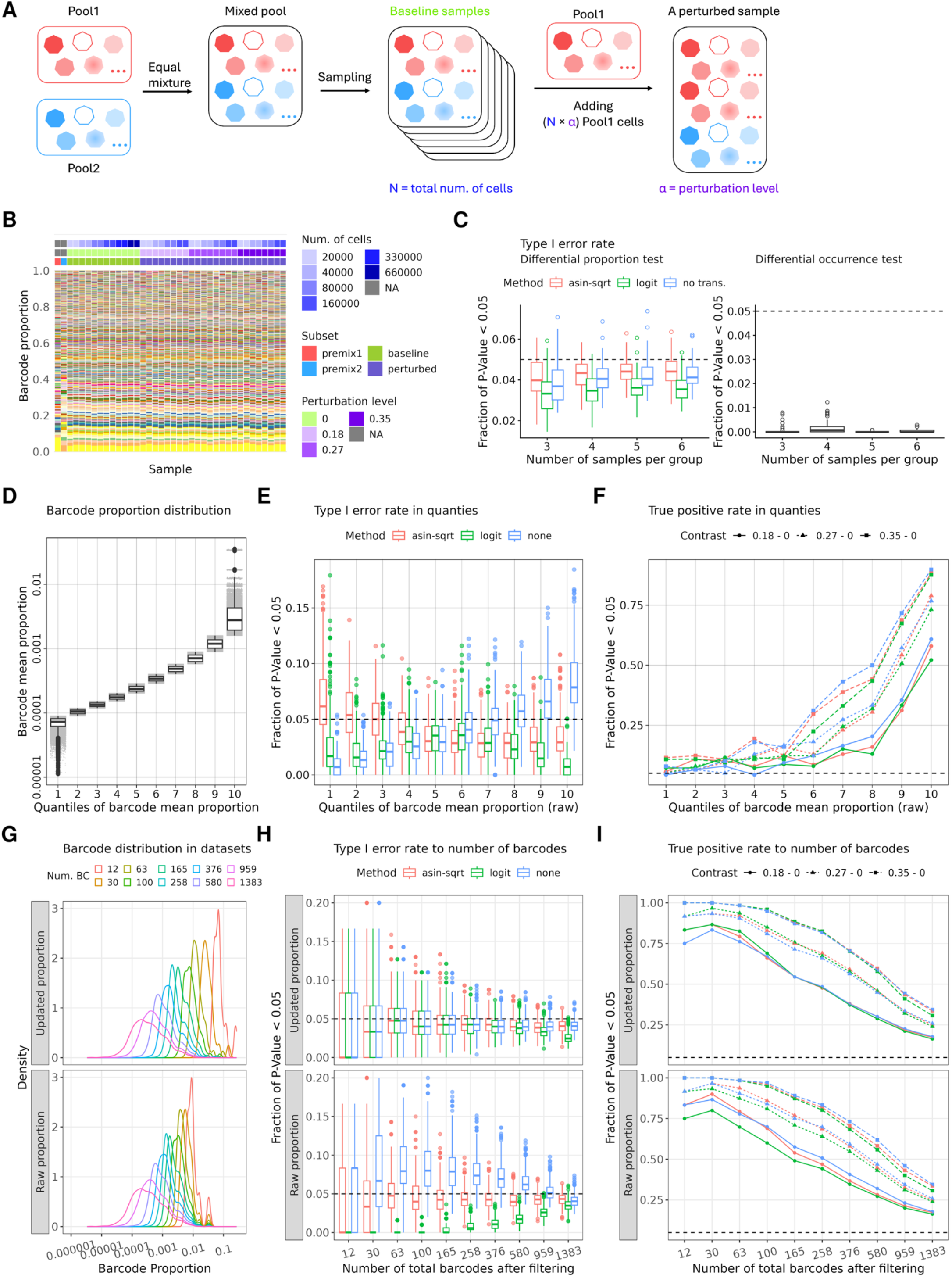
Performance of *barbieQ* statistical testing. (**A**) **Schematic of mixture experiment** showing the generation of baseline samples and perturbed samples in the Mixture dataset; (**B**) **Barcode proportion data used in evaluations** displaying proportion per sample; Samples are annotated by total number of cells and perturbation levels; (**C**) **Type** I **error rates** of (**left**) differential proportion test with three transformation methods and (**right**) differential occurrence test, taking the faction of barcodes with raw *P*-value < 0.05 obtained from each simulation (100 simulations per group size); (**D**) **Barcode proportion quantiles:** boxplot of barcode mean proportion across baseline samples in each group, binned into 10 quantiles of the mean proportion; (**E**) **Type** I **error rate as a function of barcode quantiles:** taking the fraction of barcodes with *P*-value < 0.05 within each quantile bin from Figure 3**D** using differential proportion test with three transformations in baseline samples; (**F**) As in **E** but showing power as determined by the fraction of p-values<0.05 in perturbed samples; (**G**) **Density of barcode proportion** in datasets generated by isolating different sets of barcodes, (**top**) with proportion updated per sample or (**bottom**) keeping raw proportion; datasets color-coded by number of total barcodes in datasets; (**H**) **Type** I **error rate as a function of total barcode number:** taking the faction of barcodes with *P*-value < 0.05 obtained from testing each set of barcodes in Figure 3**F** using differential proportion test with three transformations from baseline samples; (**I**) As in **H** but showing power as determined by the fraction of p-values <0.05 in perturbed samples.

Our filtering retained 1383 barcodes from the total 3998 (Supplementary Figure1). To investigate how transformation influences the differential proportion test, we applied the test after the arcsine square root (asin-sqrt), logit and no transformation. We found that the differential proportion test, across all transformations controlled the median Type I error rate below 0.05 across 100 simulations (Figure 3C). The differential occurrence test also held its size (Figure 3C). We further binned barcodes into 10 quantiles based on their proportion (Figure 3D), with the asin-sqrt transformation showing the most consistent performance across bins while logit transformation resulted in a higher Type 1 error for lower proportions and no transformation had the reverse trend (Figure 3E).

We also assessed the differential proportion test when the number of barcodes in an experiment varied, noting that barcode proportions generally decrease as the number of barcodes increase. To do thise generated datasets with differing total numbers of barcodes (N = 12, 30, 63, 100, 165, 258, 376, 580, 959, 1383) by applying various filtering thresholds (0.1, 0.2, 0.3, 0.4, 0.5, 0.6, 0.7, 0.8, 0.9, 0.95) in the `tagTopBarcode` function to the Mixture data, and extracting the top barcodes (Figure 3G). We re-calculated the barcode proportion within each dataset and repeated the Type I error rate assessment as previously described. The three transformations showed similar performance and the Type I error rate became more stable towards 0.05 as barcode size increased (Figure 3H). In addition, we assessed the Type I error rate of the differential proportion test as a function of the number of barcodes by using the raw barcode proportion for various sets of barcodes from the Mixture data. The raw proportion is generally smaller than the updated proportion (Figure 3G). Of the three transformations, asin-sqrt showed the most consistent Type I error rate control across barcode number (Figure 3H).

Finally, we evaluated Type I error rate in two further datasets (Supplementary Figure 2). For the HSPC xenograft data, samples collected from the same donor (C21, C22 and C23) were treated as null samples, despite the differences in time points, tissues, and cell types. For the AML data, samples treated by DMSO were taken as null samples, despite being collected at different times. All tests maintained Type I error rates under 0.05. A similar trend in terms of the effect of transformations was also observed in these data.

#### Power to detect true positives in real data

Using a second subset of samples in the preprocessed Mixture data, we evaluated the power of the differential proportion test to detect true positives. Twenty-four perturbed samples were generated by introducing additional Pool1 cells into the baseline level: sampling a certain number of cells from the mixed pool (n= 20, 40, 80 or 160 thousand) and adding a certain ratio of cells from Pool1 (α = 0.18, 0.27, or 0.35), with 2 replicates in each case. Specifically, (n × α) cells from Pool1 were added to the n cells from the mixed pool to generate a perturbed sample. The 12 baseline samples were considered as perturbation level α = 0. (Figure 3A) This design results in a known difference in barcode proportions (for all barcodes) between samples with different α.

We gauged the power of the differential proportion test based on the fraction of barcodes with *P*-values < 0.05, obtained from testing three contrasts using the `“testBarcodeSignif”` function: group_α=0.18_, group_α=0.27_, and group_α=0.35_ v.s. group_α=0_, respectively, each with asin-sqrt, logit and no transformation (see methods). To assess the power at different proportion ranges, we binned barcodes into 10 quantiles based on their proportion as previously described. Across all transformations, the differential proportion test tended to detect more significance in quantile bins of higher proportion or under stronger contrast, but logit generally showed lower power (Figure 3F) except at the smallest proportions.

As before, we also assessed the power of the differential proportion test under scenarios of different numbers of barcodes. We generated datasets by selecting various numbers of top barcodes from the Mixture data as previously described, and likewise, re-calculated barcode proportion per sample and repeated the power assessment for each dataset. The three transformations showed similar power, with all demonstrating greater power with lower number of barcodes and greater proportions (Figure 3I). To investigate the impact of the way proportion is calculated, we repeated this process by using the raw barcode proportion when taking subsets of barcodes at each barcode size and found asin-sqrt showed greater power than logit across barcode sizes (Figure 3I). All tests showed greater power under stronger contrast.

The coefficients of the contrasts obtained from the `“testBarcodeSignif”` function are the estimated differences between sample groups for each barcode which can be predicted from the design (see methods). Both asin-sqrt and no transformation accurately estimated the barcode proportion change, while logit tended to over-estimate the change (Supplementary Figure 3A). Residuals from asin-sqrt-transformed proportion were closest to Normality and exhibited the most consistent variance across proportion levels (Supplementary Figure 3B, C), in line with the assumption of residual Normality and homoscedasticity for the linear regression model used in the differential proportion test.

### Case study

#### Reanalysis of monkey HSPC data

Two important features of the *barbieQ* package are its data-driven filtering strategy and robust statistical tests. To demonstrate how *barbieQ* advances upon previous approaches in the clonal tracking and lineage tracing fields, we applied the *barbieQ* analytical workflow to a real-world dataset and compared the results with those reported in the original publication. The example data is a subset of 30 samples from the monkey HSPC dataset, collected across 6 time points and sorted into 5 different cell types (Figure 4A). In total, 16,603 unique barcodes were observed in this dataset, with barcode proportions ranging from a maximum of 0.14, a mean of approximately 6×10^−5^ and a median of 0 (Figure 4B). These barcode proportions are variable between cell types but relatively stable between time points within the same cell type (Figure 4B).

**Figure 4.**
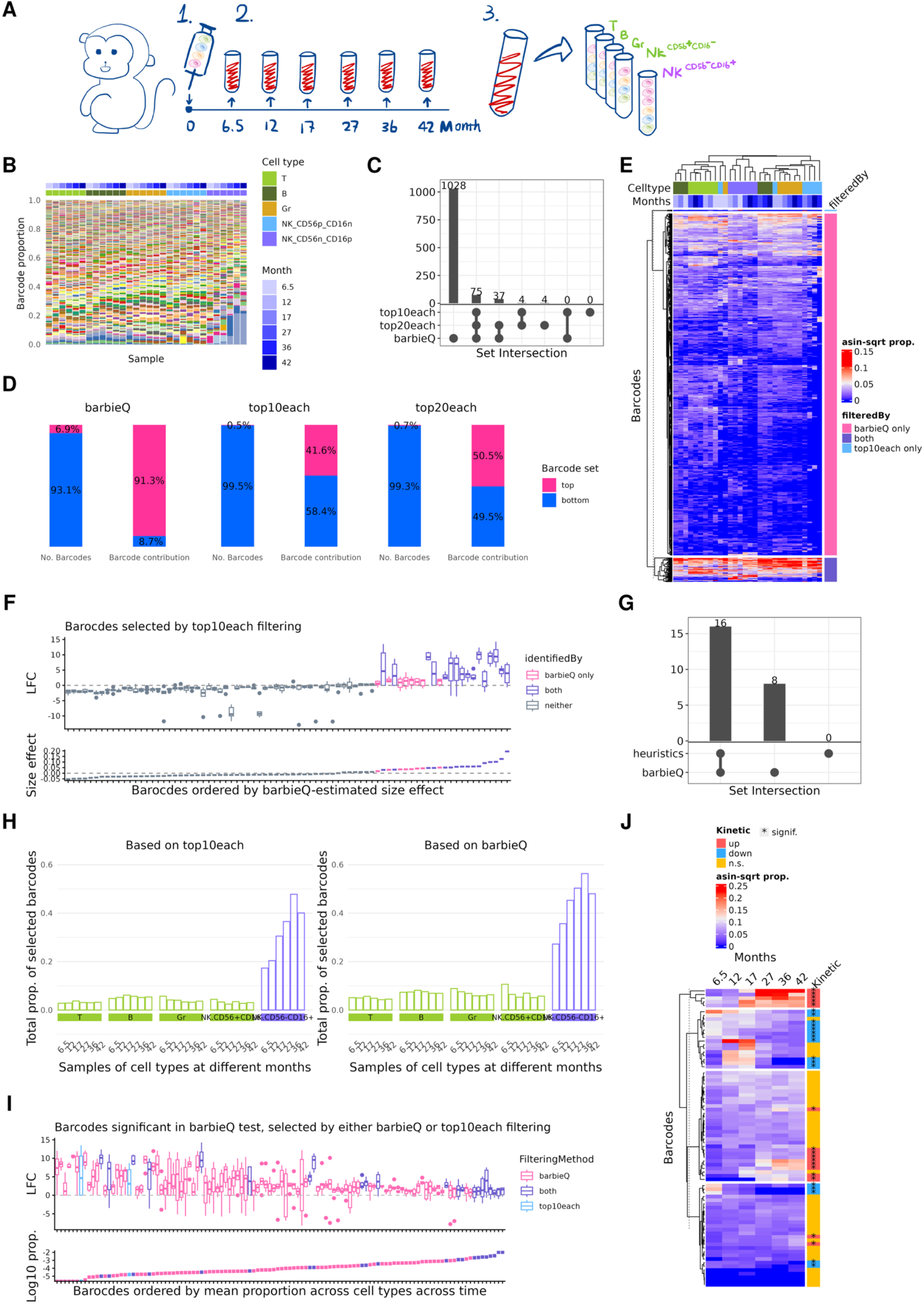
Applying *barbieQ* to the monkey HSPC dataset. (**A**) Experimental design for a subset of the monkey HSPC dataset, where 30 samples contained 5 cell types sharing 6 time points; (**B**) Barcode proportion per sample in the subset Monkey HSPC dataset, with each barcode represented by a unique colour; Samples are annotated by cell type and time to collection in months; (**C**) UpSet plot showing the overlap in retained barcodes using *barbieQ* filtering, *top10-per-sample* and *top20-per-sample* strategies; (**D**) Faction of retained barcodes and fraction of their total proportion to the whole dataset. Filtering strategies from left to right: *barbieQ, top10-per-sample*, and *top20-per-sample*; (**E**) Heatmap of retained barcodes by either *barbieQ* or *top10-per-sample*. Right annotations indicate the overlap between the two approaches; (**F**) Box plot of the LFC for each barcode. A dot refers to the LFC in proportion between to NK^CD56-CD16+^ and other cell types at a given time points (6 time points in total). Barcode were ordered by *barbieQ*-estimated coefficient for the contrast, from low to high. Barcodes were categorized into identified as NK^CD56-CD16+^-associated by *barbieQ*, heuristic LFC, both and neither method. (**G**) UpSet plot showing the overlap in identified NK^CD56-CD16+^-associated barcodes using *barbieQ* `“testBarcodeSignif”` based on *barbieQ* filtering and *top10-per-sample* strategies. (**H**) Total barcode proportion per sample for identified NK^CD56-CD16+^-associated barcodes based on *top10-per-sample* (left) and *barbieQ* filtering strategy (right), respectively. (**I**) Box plot of the LFC for each barcode. A dot refers to the LFC in proportion between to NK^CD56-CD16+^ and other cell types at a given time point (6 time points in total). Barcodes were ordered by mean proportion across cell types and time points, from low to high. Barcodes were categorized into identified as NK^CD56-CD16+^-associated by *barbieQ*, based on the set of *barbieQ* filtering and top10-per-sample, and both strategies. (**J**) Heatmap of 39 barcodes retained from 8 NK^CD56-CD16+^ samples using *barbieQ* filtering. Samples were order over time. Right annotations indicate barcodes being identified with significantly increasing, decreasing kinetics or not-significant (n.s.) over time. Barcodes were grouped into 5 patterns using the left dendrogram.

We preprocessed the raw barcode count data following the steps outlined in Figure 2, including aggregating highly correlated barcodes and filtering low-count barcodes. Compared to the “topN” filtering approaches used in the original study, *barbieQ* retained a larger number of barcodes (Figure 4C). *barbieQ* retained 8% of barcodes, accounting for 93% of the total counts while *top10-per-sample* retained 0.5% of barcodes contributing 43% of total counts, and *top20-per-sample* retained 1% of barcodes contributing 51% of total counts (Figure 4D). Figure 4E shows a heatmap of barcode logCPM, annotating barcodes retained by both *barbieQ* and *top10-per-sample* strategies or exclusively by one. Notably, barcodes exclusively retained by *barbieQ* still show informative logCPM across samples. The 4 barcodes retained by *top10-per-sample* only showed a contribution in only few samples. An additional advantage of *barbieQ* is avoiding arbitrarily selecting a *topN* threshold for every new dataset, making the filtering process less subjective.

Next, we applied the statistical testing methods implemented in *barbieQ* by reproducing the comparisons from the original publication using the exact same sets of barcodes. The first analysis included 81 barcodes that were selected by the *top10-per-sample* filtering strategy out of the 30 samples (Figure 4A, B). We used *barbieQ* to test barcode differences between the NK^CD56-CD16+^ samples and other cell types (T, B, Gr and NK^CD56+CD16-^), treating each individual time point as a factor. We used a design matrix that sets the 5 cell types as 5 factors and 6 time points as 6 additional factors with the contrast as NK^CD56-CD16+^ -(T + B + Gr + NK^CD56+CD16-^) / 4. We then applied the differential proportion test, using the `“testBarcodeSignif”` function, with the asin-sqrt transformation. We identified 23 barcodes with significantly increased proportion in NK^CD56-CD16+^ samples, including 6 novel barcodes and 17 overlapping with those selected by the original publication, which used a heuristic cutoff for barcode proportion ratios between NK^CD56-CD16+^ and other cell types at each time point respectively (Figure 4F). We inspected the log fold change (LFC) in barcode proportions (NK^CD56-CD16+^ / other cell types) at different time points and observed that the 6 novel barcodes showed robust LFC although with lower effect size (Figure 4F).

We also applied the same test to a larger set of barcodes retained using the *barbieQ* filtering strategy, including 1,028 barcodes that contribute 91.3% to the 30 samples. We identified three times more barcodes showing significantly increased proportion in NK^CD56-CD16+ cells^ and collectively contributing more to the NK^CD56-CD16+^ samples (Figure 4G, H). We inspected the LFC of the significant barcodes obtained based on each filtering strategy. (Figure 4J) Using the *barbieQ* filtering strategy allowed us to keep more barcodes with robust LFC and relatively high abundance.

The second analysis used the subset of 8 NK^CD56-CD16+^ samples from the monkey HSPC dataset, collected at various time points. We used the 39 barcodes retained using the *top10-per-sample* filtering strategy to match the set of barcodes in the original publication. We then used the `“testBarcodeSignif”` function to test barcodes that change over time within NK^CD56-CD16+^ samples. Specifically, each sample was given a numeric factor for the number of months post transplantation. We identified 17 barcodes with significantly increased and 8 with decreased proportions over time (Figure 4I). The *barbieQ* testing results robustly categorized barcode kinetics, compared to the barcodes selected by visual inspection of heatmap dendrograms as used in the original publication.

## Discussion

Lineage tracing experiments provide powerful insights into cellular development and evolution, but the complexity of the data presents significant analytical challenges. Current approaches, typically based on pairwise comparisons, may fail to account for the inherent complexity of the experimental design, potentially leaving important insights unexplored. Here, we presented *barbieQ*, a new R Bioconductor package tailored to the analysis of lineage tracing barcode count data that provides an integrated platform for quality control, filtering, visualisation and statistical testing. Built upon the *SummarizedExperiment* infrastructure, it ensures compatibility with existing workflows. *barbieQ*’s core function `testBarcodeSignif` provides statistical tests for identifying differential barcode proportions and differential barcode occurrence. These tests enable the robust identification of barcode differences between sample groups, whilst comprehensively accounting for variability resulting from multiple technical and biological conditions. *barbieQ*’s data-driven approach to preprocessing using the functions `clusterCorrelatingBarcodes` and `tagTopBarcodes`, guarantees a clean, reproducible and informative set of barcodes for testing. Additionally, the built-in visualisation functions can be used to inspect barcode features throughout an analysis, ensuring transparency and interpretability at every stage of the workflow.

We demonstrated *barbieQ*’s preprocessing capabilities by applying the `clusterCorrelatingBarcodes` function to four barcode datasets: monkey HSPC, AML data, HSPC xenograft data, and mixture data. In the first three, we found highly correlated barcode clusters likely to be the multiple barcodes that were incorporated into the same initial cells during preparation. We previously applied this approach in the analysis of an unpublished dataset and identified clusters of highly correlated barcodes, which were also detected in the same initial cells by single cell sequencing. Although this approach was validated as useful for confirming barcode uniqueness, its reliance on heuristic thresholds may limit generalisation across different datasets. We also applied the `tagTopBarcodes` function to the four datasets to identify the main contributing barcodes across samples. We showed that our data-driven method effectively filters the data whilst retaining the informative barcodes. In the case study, we compared the *barbieQ* filtering strategy to the commonly used *top-10* (or *20*)-per-sample approach in the analysis of Monkey HSPC data, where our approach retained a greater number of potentially essential barcodes.

Using the Mixture data, we evaluated the *barbieQ*’s statistical tests by examining Type I error rate by simulating sample groups under null conditions and the power by detecting known barcode differences using the function `testBarcodeSignif`. We showed that both statistical tests, differential proportion and differential occurrence, held their sizes. The differential proportion test combines *limma* with commonly-used transformation methods for proportion data; the asin-sqrt and logit transformations, in order to accommodate the assumptions of normally distributed data and mean variance relationship in the linear regression model.^50^ Based on our assessment, we recommend using asin-sqrt in barcode data analysis for its superior performance. Specifically, we found asin-sqrt transformation had: 1) consistent Type I error rate across proportion levels, across datasets with different numbers of barcodes and with or without updating proportion within selected barcodes, 2) greater power to detect true positives, and 3) improved residual Normality and reduced heteroscedasticity.

Both *barbieQ* statistical tests are regression-based, and their performance depends on how well the underlying assumptions are satisfied. In extreme cases, e.g., strong outliers, very high sparsity or small sample size, non-parametric approaches may be preferable, though we have not encountered such scenarios in the datasets analysed here. Since the *barbieQ* differential proportion test leverages *limma*, readers may wonder whether classic differential gene expression (DE) tools such as edgeR are suitable for barcode data. Unlike gene expression, where negative binomial (NB) models capture gene-specific baseline abundance and dispersion, barcode counts reflect clonal abundance driven primarily by biological conditions rather than clone-specific baseline levels.^45,54^ *barbieQ* therefore adopts a proportion-based framework, offering a more principled and interpretable approach to clonal composition analysis. NB-based methods may perform adequately in practice, but their assumptions are less well-aligned with barcode data.

We also applied the *barbieQ* workflow to a case study analysis of the monkey HSPC data, in which samples differed across time points and cell types. We repeated analyses from the original paper using the *barbieQ* differential proportion test and identified a novel set of barcodes with significantly higher proportion in mature NK cells compared with other cell types. We also robustly identified a set of barcodes showing significantly increasing proportion over time in the mature NK cell cohort, some of which were only described using a heatmap in the original paper. This demonstrates that *barbieQ* can recapitulate existing results derived using heuristics and visualisation, whilst robustly and reproducibly identifying additional barcodes of interest. Furthermore, given that *barbieQ*’s differential proportion test is built upon the versatile *limma* framework, its usage can be extended beyond simple group comparisons, such as fitting “splines” to identify more complex barcode kinetics.

Beyond the analyses demonstrated here, it is worth situating barbieQ within the broader landscape of genetic barcoding methods. Such studies broadly follow two analytical directions: lineage tree reconstruction and clonal analysis. Tools such as CoSpar^35^ infer lineage relationships by jointly modelling clonal and transcriptomic information, whereas barbieQ focuses on clonal analysis, statistically identifying clones with significant changes in abundance or occurrence. These approaches are complementary, and integration with lineage inference tools represents a promising direction for future development of barbieQ.

In conclusion, *barbieQ* is a powerful R Bioconductor package for robustly and reproducibly detecting biologically meaningful changes in clonal analysis using barcode count data. The *barbieQ* package is available from Bioconductor.^46^

## Methods

### Software and code availability

The *barbieQ* package was developed using R 4.5.0 and Bioconductor (version 3.21) and is available at https://www.bioconductor.org/packages/release/bioc/html/barbieQ.html. A package vignette is provided to guide users through a typical workflow using an example dataset and is available at https://www.bioconductor.org/packages/release/bioc/vignettes/barbieQ/inst/doc/barbieQ.html.

To ensure high performance and interoperability within the R Bioconductor ecosystem we leveraged established libraries such as *Dplyr* (https://cran.r-project.org/web/packages/dplyr/index.html) version 1.1.4 for data manipulation, *tydyr* (https://cran.r-project.org/web/packages/tidyr/index.html) version 1.3.1 for numerical calculations, *SummarizedExperiment* (https://bioconductor.org/packages/release/bioc/html/SummarizedExperiment.html) version 1.37.0 and *S4Vectors* (https://bioconductor.org/packages/release/bioc/html/S4Vectors.html) version 0.45.4 for data structure, *ggplot2* (https://cran.r-project.org/web/packages/ggplot2/index.html) version 3.5.1 and *ComplexHeatmap* (https://www.bioconductor.org/packages/release/bioc/html/ComplexHeatmap.html) version 2.23.0 for drawing plots, and *stats* (https://cran.r-project.org/doc/manuals/r-patched/packages/stats/refman/stats.html) version 4.5.0, *limma* (https://bioconductor.org/packages/release/bioc/html/limma.html) version 3.63.11 and *logistif* (https://cran.r-project.org/web/packages/logistf/index.html) version 1.26.0 for statistical modelling.^48,52^

The package was structured based on the *SummarizedExperiment* S4 class, allowing data manipulation and processing directly within the data structure. Processed results are saved to back to the same structure, extended with additional features to seamlessly interoperate with upstream barcode alignment tools that also use *SummarizedExperiment* or *DGEList* from *edgeR* (https://bioconductor.org/packages/release/bioc/html/edgeR.html) for data handling.^54^

The analysis code used to generate Figures 2-4 and Supplementary Figure 1-3 has been archived on Zenodo at https://doi.org/10.5281/zenodo.20078562, with the corresponding GitHub repository also provided at https://github.com/Oshlack/barbieQ-paper-analysis.

### Data description

Several published datasets, briefly described below, were used to demonstrate preprocessing and evaluate the statistical tests implemented in *barbieQ*. Raw barcode count data were obtained following sequence-level QC and barcode library alignment, which had been completed by the original authors.

#### Monkey HSPC Data

A subset of publicly available data from a study on monkey hematopoietic stem and progenitor cell (HSPC) clonal expansion *in vivo* using barcoding technique.^55^ The monkey HSPC data were analysed using the *barcodetrackR* package and made available via the compatible barcodetrackR Data repository on GitHub (https://github.com/dunbarlabNIH/barcodetrackRData/tree/main/datasets/WuC_etal).^28^ Herein, we used data from monkey “ZG66” including 16,603 barcodes across all samples, where we further selected 30 samples to be used here.

Briefly, unique barcodes were initially integrated into the DNA of HSPCs and subsequently passed to progeny cells. At a series of time points, progeny cells were collected from blood and sorted into various cell types, including T cells, B cells, granulocytes (Gr), and natural killer (NK) cells for NK^CD56+CD16-^ and NK^CD56-CD16+^ subtypes. Barcode counts across different cell types were used to interpret the patterns of HSPC differentiation. The original study focused on identifying barcodes (clones) with higher abundance in NK^CD56-CD16+^ samples compared to other cell type samples.

#### AML Data

Publicly available data from a study of acute myeloid leukaemia (AML) clones, investigating the heterogeneity in their response to various therapeutic drugs *in vitro*, which was originally analysed and introduced with *bartools*.^31^ Briefly, AML cells were barcoded at the DNA level using the SPLINTR system, recognized as individual clones, and their population was expanded under exposure to different drugs (Arac, IBET), at various doses and DMSO as a negative control.^22^ Samples were collected from each treatment condition at a series of time points. This barcode count matrix contains 1,811 barcodes, across 41 samples, under the described conditions.

#### HSPC Xenograft Data

Public data from a study of engraftment tracking of human umbilical cord blood HSPC xenotransplanted into mice.^42^ The data were analysed in the *barcodetrackR* publication and made available via the compatible barcodetrackR Data repository on GitHub (https://github.com/dunbarlabNIH/barcodetrackRData/tree/main/datasets/BelderbosME_etal).^28^ In this study, human cord blood HSPC cells from 20 individual donors were isolated and barcoded at the DNA level respectively. Sets of starting clones from different donors (n = 20) were transplanted into eith one or two recipient mice. For each mouse, progeny cells were collected from peripheral blood at multiple time points, as well as from various tissues. From each collection, cells were sorted into different cell types forming individual samples, with unsorted cells also retained. Herein, we used data of recipient mice with sufficiently engrafted HSPC clones, containing 10,149 barcodes across 8 donors, with 199 samples, under the described conditions.

#### Mixture Data

This dataset was generated to simulate both true and null changes in barcode abundance by mixing cells from two barcoded cell line samples.^43^ Cells from each cell line were divided into two pools (Pool1 and Pool2), each incorporating distinct clonal tracking barcodes into their DNA, ensuring no overlap between pools. Cells in each pool were counted and mixed in an equal ratio to produce a mixed pool. Twelve baseline samples were sampled from the mixed pool at various cell numbers. Twenty-four perturbed samples were generated by sampling a certain number of cells from the mixed pool and adding a certain ratio of cells from Pool1, with 2 replicates in each case. This dataset contains 3998 barcodes, across 36 samples as described, as well as two reference samples representing the unmixed Pool1 and Pool2 barcode counts.

### Data analysis

All data was analysed using the functions provided in barbieQ. Complete documentation of these can be found in the analysis on GitHub (https://github.com/Oshlack/barbieQ-paper-analysis).

#### True positive calculations for Mixture data

The mixture pool contains a mix of cells with barcodes from Pool1 and Pool2, for which the barcodes didn’t overlap. Let x_n_ be the proportion of barcode, n = (1, …, 1553).

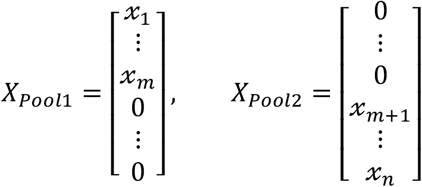

A baseline sample theoretically consisted of an equal mixture of total barcode numbers from Pool1 and Pool2, leading to barcode proportions being half of the reference samples.

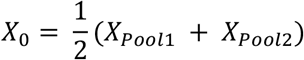

A sample with a perturbation level of α theoretically leads to a known change in proportion for the barcodes from both Pool1 and Pool2.

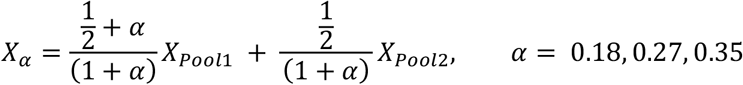

The designed difference in barcode proportions between perturbed samples and baseline samples can therefore be calculated for each transformation.

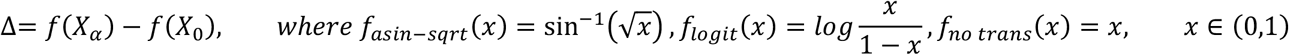

Therefore, for this data we modelled the 36 samples using a design matrix that set factors for each perturbation level (α = 0, 0.18, 0.27, and 0.35), with additional factors accounting for the cell number in the equal mixture (n= 20, 40, 80, 160, 330, and 660 thousand). We then applied the `“testBarcodeSignif”` function to test barcode differential proportion under three contrasts: group_α=0.18_, group_α=0.27_, and group_α=0.35_ v.s. group_α=0_, respectively, each with asin-sqrt, logit and no transformation. After preprocessing, a total of 1553 barcodes were tested using their raw proportion, and all barcodes were designed to change under each contrast.

## Acknowledgements

We would like to thank our collaborators, Mark Dawson, Elanor Wainright, Ali Motazedian, Enid Lam, Dane Vassiliadis, Henrietta Holze, who brought in valuable data, interesting biology questions and inspiring discussion. We would like to thank our colleagues and friends, Caitlin Page who gave advice on the package development, and Lizhong Chen who helped improve code efficiency. And we acknowledge the clonal tracking community, the discussion with whom benefited in a thorough understanding of the background and potential application for this research.

## Funding

This work has been supported by National Health and Medical Research Council [GNT1196256, GNT2026098].

### Conflict of Interest

none declared.

## Notes

### Competing Interest Statement

The authors have declared no competing interest.

### Summary of Updates

I the revised manuscript, we have focused on: 1. Clarifying the methodological scope and assumptions of the statistical tests implemented in barbieQ, including explicit statements of the null hypotheses, appropriate experimental designs, barcode structure assumptions, and the conditions under which the linear model is or is not preferable. 2. Improving the description of the preprocessing and filtering framework, including clarifications on its role, performance (sensitivity to thresholds), assumptions, and limitations. In addition, we have: 1. Replaced Figure 2C with a more informative visualization (scatter plots of barcode proportions within clusters). 2. Incorporated and discussed relevant prior work. 3. Archived the full analysis code on Zenodo and added the DOI to the manuscript. 4. Revised figures, text clarity, and documentation throughout.

## References

(1) Bramlett, C.; Jiang, D.; Nogalska, A.; Eerdeng, J.; Contreras, J.; Lu, R. Clonal Tracking Using Embedded Viral Barcoding and High-Throughput Sequencing. Nat. Protoc. 2020, 15 (4), 1436–1458. 10.1038/s41596-019-0290-z.

(2) Emert, B. L.; Cote, C. J.; Torre, E. A.; Dardani, I. P.; Jiang, C. L.; Jain, N.; Shaffer, S. M.; Raj, A. Variability within Rare Cell States Enables Multiple Paths toward Drug Resistance. Nat. Biotechnol. 2021, 39 (7), 865–876. 10.1038/s41587-021-00837-3.

(3) Gutierrez, C.; Al’Khafaji, A. M.; Brenner, E.; Johnson, K. E.; Gohil, S. H.; Lin, Z.; Knisbacher, B. A.; Durrett, R. E.; Li, S.; Parvin, S.; Biran, A.; Zhang, W.; Rassenti, L.; Kipps, T. J.; Livak, K. J.; Neuberg, D.; Letai, A.; Getz, G.; Wu, C. J.; Brock, A. Multifunctional Barcoding with ClonMapper Enables High-Resolution Study of Clonal Dynamics during Tumor Evolution and Treatment. Nat. Cancer 2021, 2 (7), 758–772. 10.1038/s43018-021-00222-8.

(4) Naik, S. H.; Perié, L.; Swart, E.; Gerlach, C.; van Rooij, N.; de Boer, R. J.; Schumacher, T. N. Diverse and Heritable Lineage Imprinting of Early Haematopoietic Progenitors. Nature 2013, 496 (7444), 229–232. 10.1038/nature12013.

(5) Lin, D. S.; Zhang, S.; Schreuder, J.; Tran, J.; Sargeant, T.; Metcalf, D.; Ng, A. P.; Weber, T. S.; Naik, S. H. A Multi-Track Landscape of Haematopoiesis Informed by Cellular Barcoding and Agent-Based Modelling. bioRxiv March 30, 2024, p 2024.03.28.587126. 10.1101/2024.03.28.587126.

(6) Chan, M. M.; Smith, Z. D.; Grosswendt, S.; Kretzmer, H.; Norman, T. M.; Adamson, B.; Jost, M.; Quinn, J. J.; Yang, D.; Jones, M. G.; Khodaverdian, A.; Yosef, N.; Meissner, A.; Weissman, J. S. Molecular Recording of Mammalian Embryogenesis. Nature 2019, 570 (7759), 77–82. 10.1038/s41586-019-1184-5.

(7) Bandler, R. C.; Vitali, I.; Delgado, R. N.; Ho, M. C.; Dvoretskova, E.; Ibarra Molinas, J. S.; Frazel, P. W.; Mohammadkhani, M.; Machold, R.; Maedler, S.; Liddelow, S. A.; Nowakowski, T. J.; Fishell, G.; Mayer, C. Single-Cell Delineation of Lineage and Genetic Identity in the Mouse Brain. Nature 2022, 601 (7893), 404–409. 10.1038/s41586-021-04237-0.

(8) Biddy, B. A.; Kong, W.; Kamimoto, K.; Guo, C.; Waye, S. E.; Sun, T.; Morris, S. A. Single-Cell Mapping of Lineage and Identity in Direct Reprogramming. Nature 2018, 564 (7735), 219–224. 10.1038/s41586-018-0744-4.

(9) Lin, D. S.; Tian, L.; Tomei, S.; Amann-Zalcenstein, D.; Baldwin, T. M.; Weber, T. S.; Schreuder, J.; Stonehouse, O. J.; Rautela, J.; Huntington, N. D.; Taoudi, S.; Ritchie, M. E.; Hodgkin, P. D.; Ng, A. P.; Nutt, S. L.; Naik, S. H. Single-Cell Analyses Reveal the Clonal and Molecular Aetiology of Flt3L-Induced Emergency Dendritic Cell Development. Nat. Cell Biol. 2021, 23 (3), 219–231. 10.1038/s41556-021-00636-7.

(10) Li, L.; Bowling, S.; McGeary, S. E.; Yu, Q.; Lemke, B.; Alcedo, K.; Jia, Y.; Liu, X.; Ferreira, M.; Klein, M.; Wang, S.-W.; Camargo, F. D. A Mouse Model with High Clonal Barcode Diversity for Joint Lineage, Transcriptomic, and Epigenomic Profiling in Single Cells. Cell 2023, 186 (23), 5183–5199.e22. 10.1016/j.cell.2023.09.019.

(11) Laisné, M.; Lupien, M.; Vallot, C. Epigenomic Heterogeneity as a Source of Tumour Evolution. Nat. Rev. Cancer 2025, 25 (1), 7–26. 10.1038/s41568-024-00757-9.

(12) VanHorn, S.; Morris, S. A. Next-Generation Lineage Tracing and Fate Mapping to Interrogate Development. Dev. Cell 2021, 56 (1), 7–21. 10.1016/j.devcel.2020.10.021.

(13) Chen, C.; Liao, Y.; Peng, G. Connecting Past and Present: Single-Cell Lineage Tracing. Protein Cell 2022, 13 (11), 790–807. 10.1007/s13238-022-00913-7.

(14) Deng, L.-H.; Li, M.-Z.; Huang, X.-J.; Zhao, X.-Y. Single-Cell Lineage Tracing Techniques in Hematology: Unraveling the Cellular Narrative. J. Transl. Med. 2025, 23 (1), 270. 10.1186/s12967-025-06318-4.

(15) Sun, J.; Ramos, A.; Chapman, B.; Johnnidis, J. B.; Le, L.; Ho, Y.-J.; Klein, A.; Hofmann, O.; Camargo, F. D. Clonal Dynamics of Native Haematopoiesis. Nature 2014, 514 (7522), 322–327. 10.1038/nature13824.

(16) Wagner, D. E.; Weinreb, C.; Collins, Z. M.; Briggs, J. A.; Megason, S. G.; Klein, A. M. Single-Cell Mapping of Gene Expression Landscapes and Lineage in the Zebrafish Embryo. Science 2018, 360 (6392), 981–987. 10.1126/science.aar4362.

(17) Spanjaard, B.; Hu, B.; Mitic, N.; Olivares-Chauvet, P.; Janjuha, S.; Ninov, N.; Junker, J. P. Simultaneous Lineage Tracing and Cell-Type Identification Using CRISPR–Cas9-Induced Genetic Scars. Nat. Biotechnol. 2018, 36 (5), 469–473. 10.1038/nbt.4124.

(18) Kong, W.; Biddy, B. A.; Kamimoto, K.; Amrute, J. M.; Butka, E. G.; Morris, S. A. CellTagging: Combinatorial Indexing to Simultaneously Map Lineage and Identity at Single-Cell Resolution. Nat. Protoc. 2020, 15 (3), 750–772. 10.1038/s41596-019-0247-2.

(19) Weinreb, C.; Rodriguez-Fraticelli, A.; Camargo, F. D.; Klein, A. M. Lineage Tracing on Transcriptional Landscapes Links State to Fate during Differentiation. Science 2020, 367 (6479), eaaw3381. 10.1126/science.aaw3381.

(20) Bowling, S.; Sritharan, D.; Osorio, F. G.; Nguyen, M.; Cheung, P.; Rodriguez-Fraticelli, A.; Patel, S.; Yuan, W.-C.; Fujiwara, Y.; Li, B. E.; Orkin, S. H.; Hormoz, S.; Camargo, F. D. An Engineered CRISPR-Cas9 Mouse Line for Simultaneous Readout of Lineage Histories and Gene Expression Profiles in Single Cells. Cell 2020, 181 (6), 1410–1422.e27. 10.1016/j.cell.2020.04.048.

(21) Oren, Y.; Tsabar, M.; Cuoco, M. S.; Amir-Zilberstein, L.; Cabanos, H. F.; Hütter, J.-C.; Hu, B.; Thakore, P. I.; Tabaka, M.; Fulco, C. P.; Colgan, W.; Cuevas, B. M.; Hurvitz, S. A.; Slamon, D. J.; Deik, A.; Pierce, K. A.; Clish, C.; Hata, A. N.; Zaganjor, E.; Lahav, G.; Politi, K.; Brugge, J. S.; Regev, A. Cycling Cancer Persister Cells Arise from Lineages with Distinct Programs. Nature 2021, 596 (7873), 576–582. 10.1038/s41586-021-03796-6.

(22) Fennell, K. A.; Vassiliadis, D.; Lam, E. Y. N.; Martelotto, L. G.; Balic, J. J.; Hollizeck, S.; Weber, T. S.; Semple, T.; Wang, Q.; Miles, D. C.; MacPherson, L.; Chan, Y.-C.; Guirguis, A. A.; Kats, L. M.; Wong, E. S.; Dawson, S.-J.; Naik, S. H.; Dawson, M. A. Non-Genetic Determinants of Malignant Clonal Fitness at Single-Cell Resolution. Nature 2022, 601 (7891), 125–131. 10.1038/s41586-021-04206-7.

(23) Goyal, Y.; Busch, G. T.; Pillai, M.; Li, J.; Boe, R. H.; Grody, E. I.; Chelvanambi, M.; Dardani, I. P.; Emert, B.; Bodkin, N.; Braun, J.; Fingerman, D.; Kaur, A.; Jain, N.; Ravindran, P. T.; Mellis, I. A.; Kiani, K.; Alicea, G. M.; Fane, M. E.; Ahmed, S. S.; Li, H.; Chen, Y.; Chai, C.; Kaster, J.; Witt, R. G.; Lazcano, R.; Ingram, D. R.; Johnson, S. B.; Wani, K.; Dunagin, M. C.; Lazar, A. J.; Weeraratna, A. T.; Wargo, J. A.; Herlyn, M.; Raj, A. Diverse Clonal Fates Emerge upon Drug Treatment of Homogeneous Cancer Cells. Nature 2023, 620 (7974), 651–659. 10.1038/s41586-023-06342-8.

(24) Weber, T. S.; Biben, C.; Miles, D. C.; Glaser, S. P.; Tomei, S.; Lin, C.-Y.; Kueh, A.; Pal, M.; Zhang, S.; Tam, P. P. L.; Taoudi, S.; Naik, S. H. LoxCode in Vivo Barcoding Reveals Epiblast Clonal Fate Bias to Fetal Organs. Cell 2025, 0 (0). 10.1016/j.cell.2025.04.026.

(25) Lin, D. S.; Kan, A.; Gao, J.; Crampin, E. J.; Hodgkin, P. D.; Naik, S. H. DiSNE Movie Visualization and Assessment of Clonal Kinetics Reveal Multiple Trajectories of Dendritic Cell Development. Cell Rep. 2018, 22 (10), 2557–2566. 10.1016/j.celrep.2018.02.046.

(26) Rodriguez-Fraticelli, A. E.; Wolock, S. L.; Weinreb, C. S.; Panero, R.; Patel, S. H.; Jankovic, M.; Sun, J.; Calogero, R. A.; Klein, A. M.; Camargo, F. D. Clonal Analysis of Lineage Fate in Native Haematopoiesis. Nature 2018, 553 (7687), 212–216. 10.1038/nature25168.

(27) Thielecke, L.; Cornils, K.; Glauche, I. genBaRcode: A Comprehensive R-Package for Genetic Barcode Analysis. Bioinformatics 2020, 36 (7), 2189–2194. 10.1093/bioinformatics/btz872.

(28) Espinoza, D. A.; Mortlock, R. D.; Koelle, S. J.; Wu, C.; Dunbar, C. E. Interrogation of Clonal Tracking Data Using barcodetrackR. Nat. Comput. Sci. 2021, 1 (4), 280–289. 10.1038/s43588-021-00057-4.

(29) Hadj Abed, L.; Tak, T.; Cosgrove, J.; Perié, L. CellDestiny: A RShiny Application for the Visualization and Analysis of Single-Cell Lineage Tracing Data. Front. Med. 2022, 9, 919345. 10.3389/fmed.2022.919345.

(30) Putri, G. H.; Pires, N.; Davidson, N. M.; Blyth, C.; Al’Khafaji, A. M.; Goel, S.; Phipson, B. Extraction and Quantification of Lineage-Tracing Barcodes with NextClone and CloneDetective. November 20, 2023. 10.1101/2023.11.19.567755.

(31) Holze, H.; Talarmain, L.; Fennell, K. A.; Lam, E. Y.; Dawson, M. A.; Vassiliadis, D. Analysis of Synthetic Cellular Barcodes in the Genome and Transcriptome with BARtab and Bartools. Cell Rep. Methods 2024, 100763. 10.1016/j.crmeth.2024.100763.

(32) Sun, W.; Perkins, M.; Huyghe, M.; Faraldo, M. M.; Fre, S.; Perié, L.; Lyne, A.-M. Extracting, Filtering and Simulating Cellular Barcodes Using CellBarcode Tools. Nat. Comput. Sci. 2024, 4 (2), 128–143. 10.1038/s43588-024-00595-7.

(33) TeamPerie/CellBarcodeSim at v1. GitHub. https://github.com/TeamPerie/CellBarcodeSim (accessed 2025-09-11).

(34) Weinreb, C.; Klein, A. M. Lineage Reconstruction from Clonal Correlations. Proc. Natl. Acad. Sci. 2020, 117 (29), 17041–17048. 10.1073/pnas.2000238117.

(35) Wang, S.-W.; Herriges, M. J.; Hurley, K.; Kotton, D. N.; Klein, A. M. CoSpar Identifies Early Cell Fate Biases from Single-Cell Transcriptomic and Lineage Information. Nat. Biotechnol. 2022, 40 (7), 1066–1074. 10.1038/s41587-022-01209-1.

(36) Weinreb, C.; Rodriguez-Fraticelli, A.; Camargo, F. D.; Klein, A. M. Lineage Tracing on Transcriptional Landscapes Links State to Fate during Differentiation. Science 2020, 367 (6479), eaaw3381. 10.1126/science.aaw3381.

(37) Fennell, K. A.; Vassiliadis, D.; Lam, E. Y. N.; Martelotto, L. G.; Balic, J. J.; Hollizeck, S.; Weber, T. S.; Semple, T.; Wang, Q.; Miles, D. C.; MacPherson, L.; Chan, Y.-C.; Guirguis, A. A.; Kats, L. M.; Wong, E. S.; Dawson, S.-J.; Naik, S. H.; Dawson, M. A. Non-Genetic Determinants of Malignant Clonal Fitness at Single-Cell Resolution. Nature 2022, 601 (7891), 125–131. 10.1038/s41586-021-04206-7.

(38) barcodetrackR. Bioconductor. http://bioconductor.org/packages/barcodetrackR/ (accessed 2025-09-11).

(39) TeamPerie/CellDestiny, 2025. https://github.com/TeamPerie/CellDestiny (accessed 2025-09-11).

(40) Phipsonlab/CloneDetective, 2023. https://github.com/phipsonlab/CloneDetective (accessed 2025-11-07).

(41) Vassiliadis, D. DaneVass/Bartools, 2025. https://github.com/DaneVass/bartools (accessed 2025-09-11).

(42) Belderbos, M. E.; Jacobs, S.; Koster, T. K.; Ausema, A.; Weersing, E.; Zwart, E.; Haan, G. de; Bystrykh, L. V. Donor-to-Donor Heterogeneity in the Clonal Dynamics of Transplanted Human Cord Blood Stem Cells in Murine Xenografts. Biol. Blood Marrow Transplant. 2020, 26 (1), 16–25. 10.1016/j.bbmt.2019.08.026.

(43) Akimov, Y.; Bulanova, D.; Timonen, S.; Wennerberg, K.; Aittokallio, T. Improved Detection of Differentially Represented DNA Barcodes for High-throughput Clonal Phenomics. Mol. Syst. Biol. 2020, 16 (3), e9195. 10.15252/msb.20199195.

(44) Love, M. I.; Huber, W.; Anders, S. Moderated Estimation of Fold Change and Dispersion for RNA-Seq Data with DESeq2. Genome Biol. 2014, 15 (12), 550. 10.1186/s13059-014-0550-8.

(45) Robinson, M. D.; McCarthy, D. J.; Smyth, G. K. edgeR: A Bioconductor Package for Differential Expression Analysis of Digital Gene Expression Data. Bioinformatics 2010, 26 (1), 139–140. 10.1093/bioinformatics/btp616.

(46) barbieQ. Bioconductor. http://bioconductor.org/packages/barbieQ/ (accessed 2025-09-11).

(47) Law, C. W.; Zeglinski, K.; Dong, X.; Alhamdoosh, M.; Smyth, G. K.; Ritchie, M. E. A Guide to Creating Design Matrices for Gene Expression Experiments. F1000Research 2020, 9, 1444. 10.12688/f1000research.27893.1.

(48) Ritchie, M. E.; Phipson, B.; Wu, D.; Hu, Y.; Law, C. W.; Shi, W.; Smyth, G. K. Limma Powers Differential Expression Analyses for RNA-Sequencing and Microarray Studies. Nucleic Acids Res. 2015, 43 (7), e47. 10.1093/nar/gkv007.

(49) Law, C. W.; Alhamdoosh, M.; Su, S.; Dong, X.; Tian, L.; Smyth, G. K.; Ritchie, M. E. RNA-Seq Analysis Is Easy as 1–2-3 with Limma, Glimma and edgeR. F1000Research December 28, 2018. 10.12688/f1000research.9005.3.

(50) Belinda Phipson; Choon Boon Sim; Enzo R Porrello; Alex W Hewitt; Joseph Powell; Alicia Oshlack. Propeller: Testing for Differences in Cell Type Proportions in Single Cell Data. Bioinformatics 2022. 10.1093/bioinformatics/btac582.

(51) Firth, D. Bias Reduction of Maximum Likelihood Estimates. Biometrika 1993, 80 (1), 27–38. 10.1093/biomet/80.1.27.

(52) Heinze, G.; Schemper, M. A Solution to the Problem of Separation in Logistic Regression. Stat. Med. 2002, 21 (16), 2409–2419. 10.1002/sim.1047.

(53) Benjamini, Y.; Hochberg, Y. Controlling the False Discovery Rate: A Practical and Powerful Approach to Multiple Testing. J. R. Stat. Soc. Ser. B Methodol. 1995, 57 (1), 289–300. 10.1111/j.2517-6161.1995.tb02031.x.

(54) Chen, Y.; Chen, L.; Lun, A. T. L.; Baldoni, P. L.; Smyth, G. K. edgeR v4: Powerful Differential Analysis of Sequencing Data with Expanded Functionality and Improved Support for Small Counts and Larger Datasets. Nucleic Acids Res. 2025, 53 (2), gkaf018. 10.1093/nar/gkaf018.

(55) Wu, C.; Espinoza, D. A.; Koelle, S. J.; Yang, D.; Truitt, L.; Schlums, H.; Lafont, B. A.; Davidson-Moncada, J. K.; Lu, R.; Kaur, A.; Hammer, Q.; Li, B.; Panch, S.; Allan, D. A.; Donahue, R. E.; Childs, R. W.; Romagnani, C.; Bryceson, Y. T.; Dunbar, C. E. Clonal Expansion and Compartmentalized Maintenance of Rhesus Macaque NK Cell Subsets. Sci. Immunol. 2018, 3 (29), eaat9781. 10.1126/sciimmunol.aat9781.

