## Supplementary for "barbieQ: An R software package for analysing barcode count data from clonal tracking experiments"

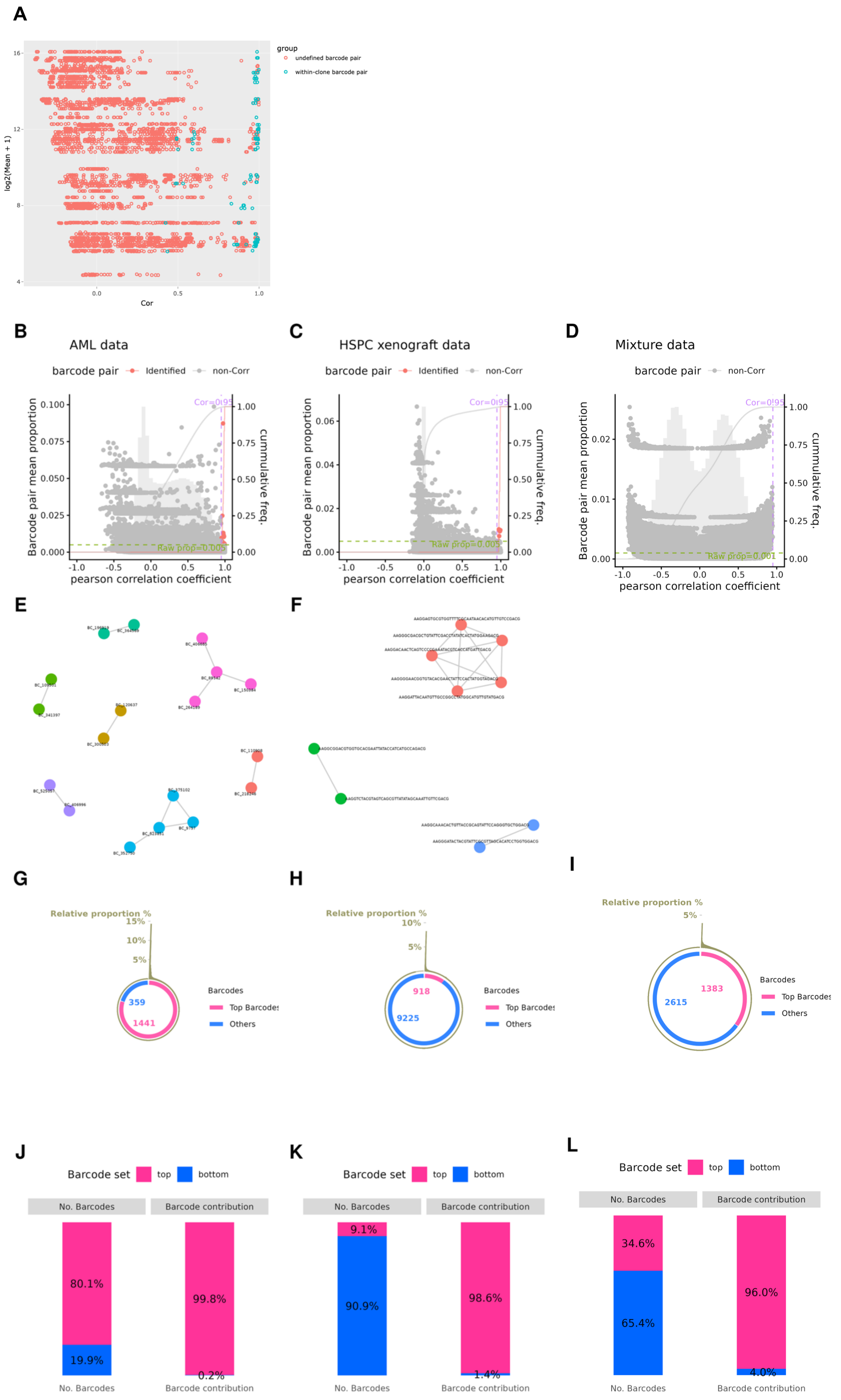
A
group
undefined barcode pair
within-clone barcode pair
log2(Mean + 1)
Cor
B
AML data
C
HSPC xenograft data
D
Mixture data
barcode pair
Identified
non-Corr
Barcode pair mean proportion
cummulative freq.
pearson correlation coefficient
Cor=0.95
Raw prop=0.005
Raw prop=0.001
E
F
G
H
I
Relative proportion %
Barcodes
Top Barcodes
Others
359
1441
918
9225
1383
2615
J
K
L
Barcode set
top
bottom
No. Barcodes
Barcode contribution
80.1%
19.9%
99.8%
0.2%
9.1%
90.9%
98.6%
1.4%
34.6%
65.4%
96.0%
4.0%

**Figure S1. Preprocessing results on an unpublished data (top row), AML data (left panel), HSPC xenograft data (middle panel), and mixture data (right panel).** (**A**) Log mean CPM and Pearson correlation coefficients for each pair of barcodes from the unpublished data (mouse haematopoietic stem cell transduced with SPLINTR lentiviral barcode library); barcode pairs are coloured by co-detection of barcodes observed in individual cells using single-cell RNA sequencing. Pairwise correlations were calculated from the proportions of 98 abundant barcodes across 222 samples collected across mice, tissues and time. (**B ~ D**) Mean proportion and Pearson correlation coefficients for each pair of barcodes; (**E, F**) Cluster of highly correlated barcodes; (**G ~ I**) Proportion of each barcode's mean CPM out of all barcodes (outside circle), color-coded by "top" or "bottom" contributors as tagged by the filtering strategy (inside circle); (**J ~ L**) Fraction of "top" or "bottom" contributor barcodes and their total contribution to the dataset.

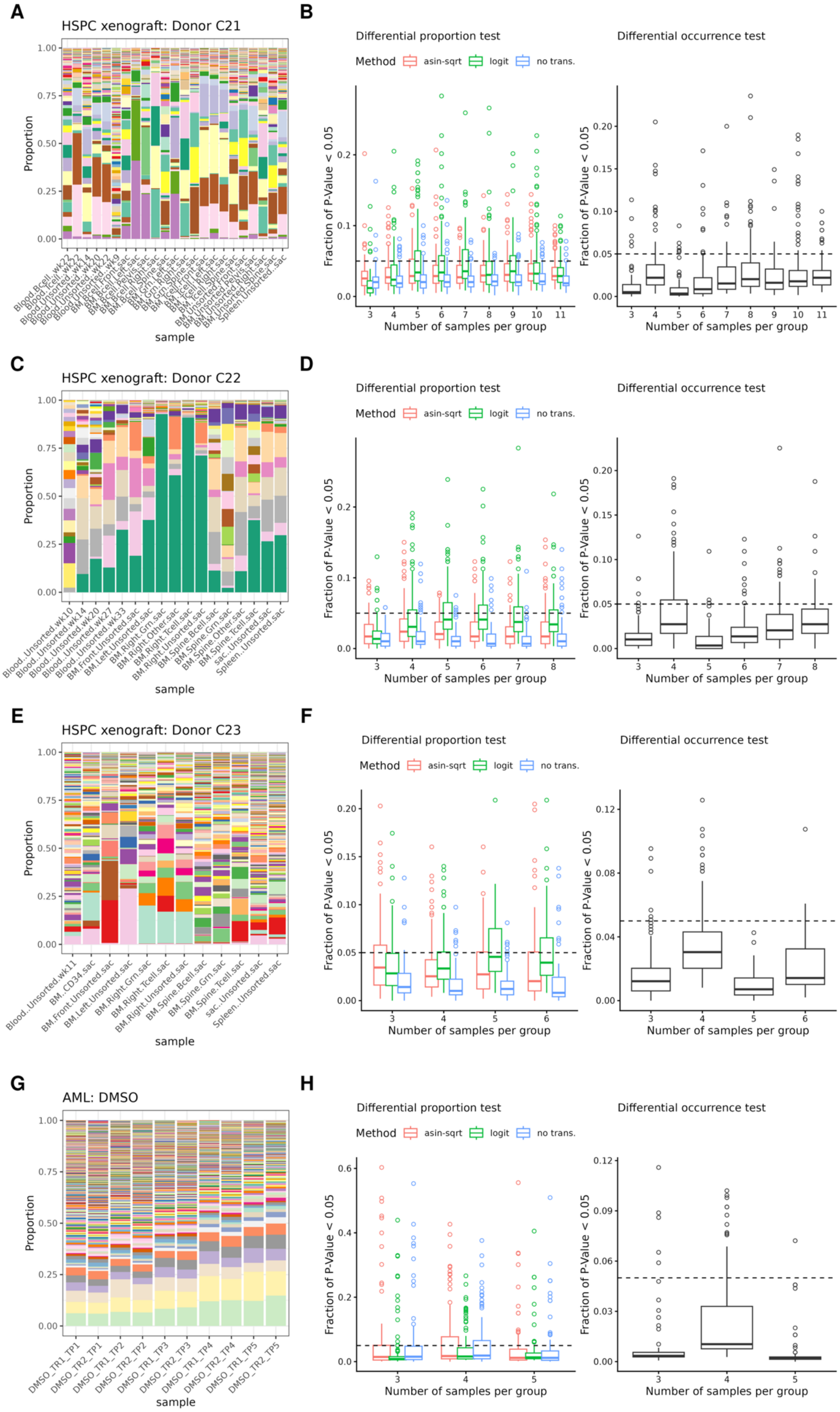


**Figure S2. Type I error rates of barbieQ statistical tests using HSPC xenograft data and AML data.** (**A, C, E, G**) Barcode proportion data used in evaluations displaying proportion per sample using samples of Donor C21, C22, and C23 from the HSPC xenograft data and samples under DMSO treatment from the AML data, with each barcode represented by a unique colour; (**B, D, F, H**) Type I error rates of (**left**) differential proportion test with three transformation methods and (**right**) differential occurrence test, taking the faction of barcodes with raw *P*-value < 0.05 obtained from each simulation (100 simulations per group size) using data in **A, C, E, G**, respectively.

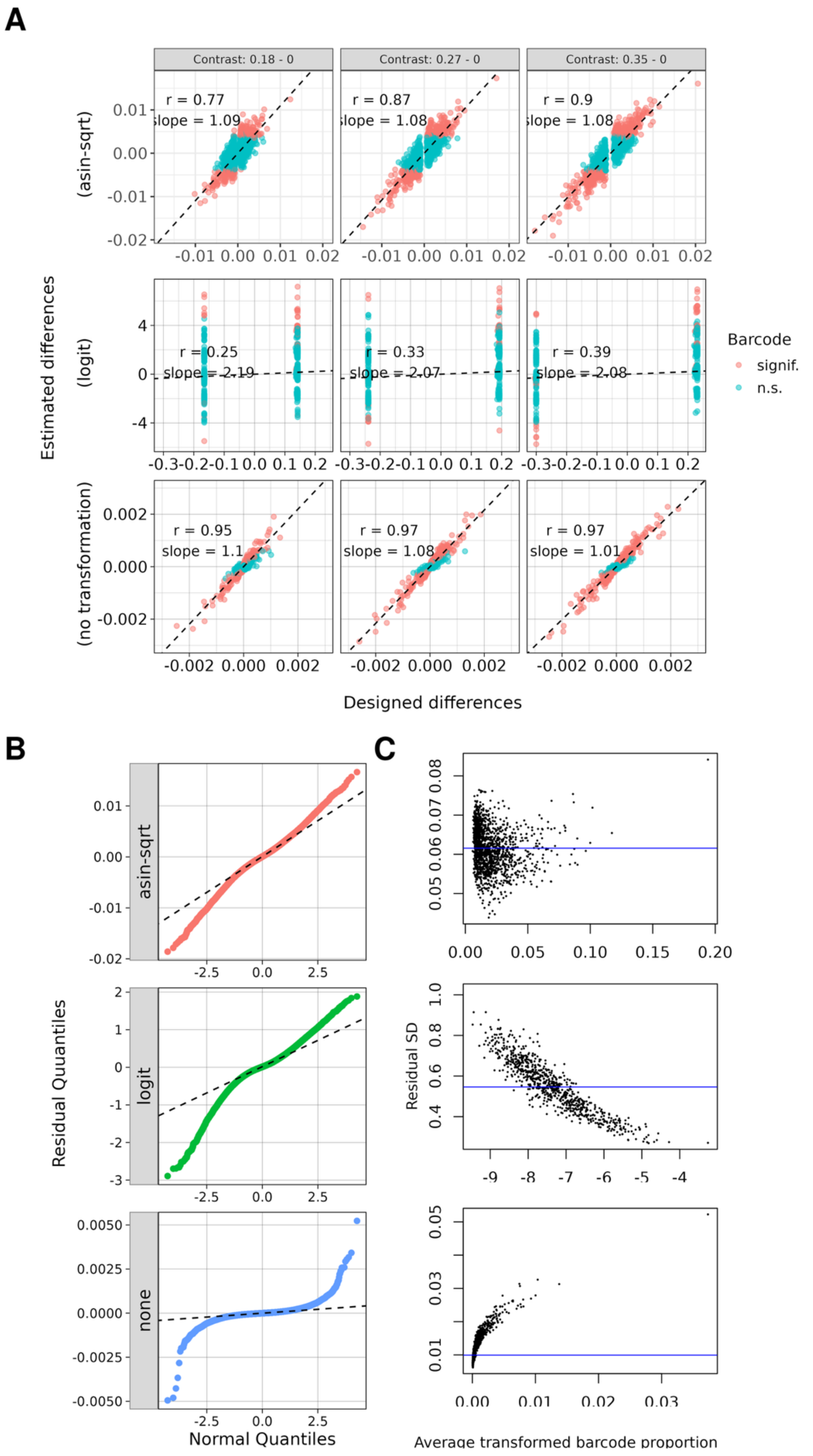


**Figure S3. Estimates of *barbieQ* differential proportion test on real contrasts.** (**A**) **Detection of true differences between samples:** estimated differences vs. designed differences. Estimated differences in barcode proportions between perturbed samples (perturbation level = 0.18, 0.27, 0.25) and baseline samples (perturbation level = 0) were obtained from the estimated coefficients of the contrast using differential proportion test with three transformation methods; Designed differences in barcode proportion for each contrast under different transformations, was calculated based on original barcode proportion in Pool1 and Pool2 samples and experimental design in Figure 3**A**; (**B**) **Normality of residuals:** QQ plots comparing residuals to a normal distribution; Residuals obtained from fitting the contrast (0.35 – 0) in (**A**) using the linear regression model implemented in the differential proportion test with three transformation methods; (**C**) **Residual variance:** standard deviation of the same residuals in (**B**), plotted against barcode mean proportion after each transformation, using *plotSA()* in *limma*.
